# Targeting NRP1 with EG00229 induces neurovascular permeability

**DOI:** 10.1101/2023.11.07.564946

**Authors:** Silvia Dragoni, James T. Brash, Alessandro Fantin, Charles Burridge, Laura Denti, Patric Turowski, Christiana Ruhrberg

**Author notes:** Department of Biosciences, University of Milan, Via G. Celoria 26, 20133 Milan, Italy. Department of Chemistry and Molecular Biology, University of Gothenburg, 405 30, Göteborg, Sweden (Charles Burridge). Francis Crick Institute, 1 Midland Rd, London NW1 1AT, UK. corresponding authors: Dr Silvia Dragoni;, Professor Christiana Ruhrberg.

## Abstract

NRP1 is a therapeutic target for inhibiting vascular endothelial growth factor (VEGF)-induced blood vessel dysfunction. The small molecule EG00229 was designed to inhibit VEGF binding to NRP1 and reduce pathological blood vessel growth. However, it is unknown whether EG00229 could also be used to reduce VEGF164-induced vascular leakage, which often exacerbates ischemic diseases due to VEGF upregulation. Here, we show that prior treatment with EG00229 prevents VEGF164-induced vascular permeability signalling, but, unexpectedly, also find that EG00229 increased rather than inhibited vascular leakage. Thus, EG00229 increased vascular leakage either when added alone or concurrently with VEGF164, both in perfused retinal explants and across primary brain EC monolayers. This EG00229-induced vascular leakage was not an off-target effect, because it relied on endothelial NRP1 expression and NRP1’s VEGF164 binding pocket, yet was independent of VEGFR1 and VEGFR2. Moreover, EG00229 activated molecular events typical of VEGF164-induced paracellular permeability, including p38 MAP kinase (p38) and SRC family kinase (SFK) phosphorylation as well as CDH5 rearrangement in endothelial junctions. Investigating EG00229-induced signalling therefore helps elucidate NRP1-dependent mechanisms of paracellular permeability induction and might help identifying new approaches to modulate the neurovascular barrier.

## Introduction

Neuropilin 1 (NRP1) acts in several cell types as a coreceptor for the VEGF165 isoform of the vascular endothelial growth factor A (VEGF) or SEMA3 proteins to variably promote normal neuronal patterning, blood vessel growth or vascular remodelling [1,2]. NRP1 also binds VEGF165 and SEMA3 proteins to drive pathological processes, for example, neovascularisation, vascular permeability, immune cell activation, tumour cell migration and metastasis [3,4,2,5–7]. Accordingly, NRP1 is a target for therapeutic developments seeking to modulate NRP1-driven processes such as vascular dysfunction, cancer progression or defective nervous system repair [4,6,7]. Thus, antibodies have been developed to target NRP1’s a1a2 versus b1 domains to selectively impair SEMA3 or VEGF165 binding to NRP1 [8,9]. Most strategies pursuing NRP1 inhibition have, however, homed in on NRP1’s binding pocket for C-end rule (CendR) peptides, whose carboxyl-terminal arginine (R)-x-x-R motif docks to a binding pocket in the NRP1 b1 domain [10]. A CendR motif is constitutively found at VEGF165’s C-terminus and is exposed at SEMA3A’s C-terminus after furin cleavage [11,12]. Furin cleavage can also expose a CendR motif in the S1 subunit of SARS-CoV-2’s spike protein, and S1 docking to NRP1 enhances viral internalisation after ACE receptor binding on target cells [13,14].

CendR-based NRP1 inhibitors include several peptides, peptidomimetics and small molecule antagonists [15–17,14,18]. When constructing peptide and peptidomimetic antagonists, the C-terminal arginine is retained for interaction with aspartate 320 (D320) in the CendR binding pocket, and in small-molecule antagonists is typically substituted by arginyl moieties that engage D320. The synthetic peptide RPARPAR also binds NRP1’s CendR pocket [11]. However, rather than acting as an inhibitor, multivalent RPARPAR, immobilised on particles, induces NRP1-mediated internalisation of payloads into cells and tissues, such as drugs to target NRP1-expressing tumour cells [11]. Moreover, multivalent RPARPAR peptides can increase vascular leakage in the Miles assay of vascular permeability and disrupt the barrier of primary human ECs (HUVEC) via NRP1’s cytoplasmic domain (NCD) in an otherwise elusive mechanism [12].

EG00229 is a small molecule that inhibits VEGF-induced, NRP1-dependent endothelial and tumour cell migration by binding NRP1’s CendR pocket [18] and has served as a prototype for other NRP1 inhibitors [17]. Prior work demonstrated that EG00229 has anti-tumoral activity. For example, EG00229 suppresses NRP1-mediated nasopharyngeal carcinoma cell migration and invasion [19]. Moreover, EG00229 increases the potency of the cytotoxic agents paclitaxel and 5-fluorouracil in lung adenocarcinoma cells [18] and synergises with lenvatinib to inhibit cholangiocarcinoma cells, which rely on NRP1 to grow and metastasise [20]. EG00229 also enhances the anti-tumour activity of microglia in a mouse glioma model [21]. With multiple roles in blocking tumour cells as well as tumour angiogenesis, EG00229 is considered useful for NRP1 blockade in cancer therapy [19]. More recently, EG00229 was reported to impair NRP1 binding of the SARS-CoV-2 S1 spike protein, which carries a CendR motif, to Caco-2 cells, thereby suggesting that NRP1 inhibition might be useful to treat COVID-19 [14]. Further, EG00229 and S1 can both compete with VEGF165 for NRP1-induced sensory neuron firing to modulate pain perception [22].

Although NRP1 inhibition has also been proposed as a therapeutic strategy to complement current anti-VEGF approaches to treat neovascular eye diseases [23–25], it has not been examined whether a CendR-based NRP1 inhibitor such as EG00229 would be suitable to reduce VEGF-induced vascular leakage from retinal vasculature. Here, we have investigated how EG00229 affects the barrier function of retinal and brain (neurovascular) endothelial cells and unexpectedly found that it increased acute vascular leakage and permeability signalling.

## Methods

### Structural models

The published crystal structures for VEGF165 in complex with NRP1 (PDB:4deq) [33] and EG00229 (chain B, PDB:3i97)[18] were rendered with PyMOL Molecular Graphics System 1.0 v2.5.0 (Schroedinger LLC) [34] and the academic version of Ligplot+ v2.2 was used to produce molecular interaction diagrams [35].

### Animals

Animal work was performed following UK Home Office Animals in Science Procedures e-Licensing (ASpeL) and institutional Animal Welfare and Ethical Review Body (AWERB) guidelines. To produce mice with an endothelial-specific NRP1, VEGFR1 (FLT1) or VEGFR2 (KDR) deletion, we combined *Cdh5*-CreER^T2^ with floxed conditional null alleles *Nrp1* (*Nrp1^fl/fl^*) [26], *Kdr* (*Kdr^fl/fl^*)[27] and *Flt1* (*Flt1^fl/fl^*) [28]. To induce gene deletion, sex-matched Cre-positive and Cre-negative young adult littermates were each injected with 0.5 mg tamoxifen (Sigma-Aldrich) in 250 µl peanut oil (Sigma) once a week for 2 weeks; the last tamoxifen injection was performed 6 days before permeability-relevant experiments. We also used NCD-deficient mice (*Nrp1^cyto/cyto^*) and mice with a CendR pocket mutation (*Nrp1^D320K/D320K^*) together with wild type littermate controls [29,30]. All mice were maintained on a C57Bl/6J background (The Jackson Laboratory). We used both male and female mice in this study, with no obvious difference in the data obtained for both sexes.

### *Ex vivo* retina permeability assay

Retinal explants were prepared as described [31]. Briefly, adult mice were culled by CO_2_ overdose and the common carotid artery was cannulated to perfuse the vasculature with heparinised saline followed by cardioplegic solution (10 mM Mg^2+^, 110 mM NaCl, 8 mM KCl, 10 mM HEPES and 1 mM CaCl_2_, pH 7.0) containing 10 µM isoproterenol, and finally with cardioplegic solution supplemented with 5 g/l Evans Blue dye and 10% bovine serum albumin (BSA) w/v (5 ml of each solution per mouse). From each mouse, both eyes were surgically removed for further microdissection. One retina was used for immunostaining (see below) and the other retina, together with the attached sclera, was flattened onto a transparent silicone base (SYLGARD 184, Merck) and maintained in position with a metal ring. These retinal explants were mounted on an upright Axiophot fluorescence microscope (Zeiss) and continuously superfused with Krebs solution (124 mM NaCl, 5 mM KCl, 2 mM MgSO_4_, 0.125 mM NaH_2_PO_4_, 22 mM NaHCO_3_ and 2 mM CaCl_2_, pH 7.4) containing 5 mM glucose and 0.1% BSA w/v. A radial vessel was cannulated using a glass needle, through which the retinal vasculature was injected with 1 mg/ml sulforhodamine B (479 Da; Merck) in Krebs solution. For permeability measurements, the fluorescent content of a microvessel was recorded continuously by time-lapse imaging using a CCD camera (Hamamatsu) and HCImageLive software (Hamamatsu) for at least 2 min. Specifically, the baseline sulforhodamine B signal was recorded for ~30 s before VEGF164 or EG00229 were added dropwise onto the retina. Recording continued for at least 90 s. Time-lapse series were analysed using ImageJ (NIH Bethesda). Pixel intensity measurements were plotted against time, and permeability values were computed by fitting data to the exponential equation Ct = C0×e−kt, whereby k = 4 P/d and d is the diameter of the vessel [32]. The difference in signal intensity of the fluorescent content of a microvessel between pre-treatment and post-treatment reflected the absolute permeability change due to treatment. Permeability measurements from at least three different *ex vivo* retinal preparations were combined and expressed as mean ± SD. Statistical analysis included a repeated measures two-way ANOVA to compare the baseline, VEGF164-induced and EG00229-induced vascular permeability in the control versus mutant retina or with and without pharmacological inhibitors in wild type retina.

### Retinal immunostaining

Mouse retinae were fixed in 4% formaldehyde in PBS for 1h and wholemount immunostained as described [31] using rabbit anti-P-p38 MAPK (Thr180/Tyr182) antibody (Cell Signaling) followed by Alexa Fluor 555–conjugated goat anti–rabbit antibodies (Invitrogen) and biotin-conjugated IB4 (Merck) followed by Alexa Fluor 488-conjugated streptavidin (Invitrogen). Images were acquired with Zeiss Axioskop 2 microscope, using a ph3 Plan-Apochromat 63x/1.40 oil objective, Hamamatsu camera and the HCImage software. Images were processed with ImageJ software version 1.52a (NIH Bethesda). Each image was acquired as an RGB colour image. Channels were split to separate the IB4 staining (green channel) from the P-p38 staining (red channel). The threshold function was used on the IB4 image to create a mask of the vascular area that was transferred onto the P-p38 image to identify vascular P-p38 staining; the P-p38 area was normalised against the IB4 area. Two-way ANOVA was used to compare genotypes and treatments.

### Pharmacological inhibitors

The NRP1 inhibitor EG00229, the VEGFR2 inhibitor SU1498, the P38 inhibitor SB203580 and the SFK inhibitor PP2 were all purchased from MERCK (Dorset, United Kingdom) and used at a concentration of 10 μM unless otherwise specified.

### Primary rat brain EC isolation, transendothelial flux assay and immunostaining

Microvessels from adult male or female rat brains were isolated by collagenase/dispase digestion and BSA/Percoll density gradient centrifugation as described [32]. Purified vessels were seeded onto 12 mm transwells (Costar) coated with collagen IV and fibronectin (Merck) and cultured in EGM2-MV media (Lonza). Media were supplemented with 5 μg/ml puromycin for the first 5 days after plating. Cells were cultured for 2–3 weeks in total until their trans-endothelial electrical resistance (TEER) plateaued at values above 200 Ω cm^2^. The TEER was measured using STX-2 chopstick electrodes connected to an EVOM voltohmmeter (World Precision Instruments). 1 mg/ml 4 kDa fluorescein (FITC)-dextran was added to the apical (‘luminal’) side of the brain microvascular ECs grown on the transwells before permeability measurements were performed as described [32]. Briefly, 50 μl samples of medium were removed from the basal chamber and replaced with fresh medium at 20 min intervals for 120 min before and after adding 50 ng/ml VEGF164 or 10 μM EG00229 to the apical or basal side of the transwell cultures. Sample fluorescence was measured with a microplate reader (Tecan Safire) and readings plotted against time. Fluorescence changes along the linear slope before and after adding an agent provided a measure of vascular permeability induction [32]. Linear slope changes of three different wells from the same cell preparation were averaged and the average considered as n = 1. Data are means ± SD of at least three independent preparations. Statistical analysis included unpaired Student’s t-test followed by a post-hoc Dunn’s test, with a significance level set at 0.05. In some experiments, primary rat brain ECs cultured on transwells were fixed in 4% formaldehyde in PBS for 15 minutes, permeabilised with acetone (−20°C) and left in blocking buffer (PBS 0.5% BSA) for 5 minutes. Cells were then immunostained using CDH5 antibody [36] diluted 1:100 in blocking buffer for 30 min followed by Alexa Fluor 555–conjugated goat anti–rabbit antibodies (ThermoFisher) diluted 1:200 in blocking buffer together with Hoechst. Images were acquired with a Zeiss Axioskop 2 microscope, using a ph3 Plan-Apochromat 63x/1.40 oil objective, Orca ER camera (Hamamatsu) and HCImage software (Hamamatsu). Images were processed with ImageJ software version 1.52a (NIH Bethesda).

### Human brain ECs

Cells from the human brain microvascular EC line hCMEC/D3 [37] were seeded on collagen I-coated 35 mm tissue culture dishes (Nunc) and grown in EGM2-MV (Lonza).

### Immunoblotting

Cell lysates from primary rat ECs or hCMEC/D3 were prepared as described [31]. Lung tissue from tamoxifen-treated mice and their littermate controls was homogenized and lysed in RIPA buffer, in the presence of 0.1% SDS, protease inhibitor cocktail 2 and phosphatase inhibitor cocktail (Sigma-Aldrich). Proteins were separated by SDS-PAGE and transferred to nitrocellulose by semi-dry electrotransfer. Membranes were blocked overnight and then incubated with the appropriate primary antibody diluted in TBS containing 0.1% Tween-20, 1% bovine serum albumin (BSA) for 2 h at room temperature; the primary antibodies used were specific for P38 (MAPK14), SRC, VEGFR2, VEGFR1 or NRP1, and for the activated (phosphorylated) forms of P38 or SRC family kinases (SFK) (1:1000; all from Cell Signaling Technology), and GADPH or tubulin (1:10000; Merck). Membranes were washed twice with TBS containing 0.1% Tween-20 and once with PBS containing 0.1% Tween-20 and then incubated with goat anti-mouse or goat anti-rabbit horseradish peroxidase (HRP)-conjugated IgG (GE Healthcare) diluted 1:10000 or 1:5000, respectively, in PBS containing 0.1% Tween-20, 1% BSA. Membranes were developed using the ECL reagents (Roche) with the Bio-Rad ChemiDoc MP Imaging System and images were acquired using the Bio-Rad Image Lab software (version 6.0.1). Protein bands were evaluated by densitometric quantification, whereby signal intensity was normalised to signal intensity from GADPH or tubulin from the same sample as a loading control. Phosphorylation levels of P38 and SKHs were normalised against the total levels of P38 or SRC. Densitometric quantification of three independent immunoblots were determined by changes in protein or phosphoprotein content normalised to tubulin or GADPH total protein loading controls, with values expressed as fold increase. Data are shown as mean ± SD. Statistical analysis included one-way ANOVA followed by post-hoc Dunnett’s tests.

## Results

### EG00229 induces retinal vascular leakage

To determine whether EG00229 functions as a competitive inhibitor in VEGF-induced vascular leakage, we used an *ex vivo* assay of mouse retina that measures the extravasation of fluorescent sulforhodamine B dye from perfused microvasculature [31], which can be recorded in real-time (**Fig. S1A**). As shown [31], 10 ng/ml VEGF164 (mouse equivalent of human VEGF165) rapidly induced dye extravasation from the retinal microvessel lumen, corresponding to a ~3-fold increase in microvessel permeability (**Fig. 1A,B**). Consistent with analogous findings on VEGF inhibition in other contexts, pre-incubating retinal explants for 15 min with 10 µM EG00229 significantly reduced VEGF164-induced dye extravasation (EG00229→VEGF164; **Fig. 1A,B**). Unexpectedly, however, adding 10 µM EG00229 alone also induced dye extravasation at a similar extent as VEGF164, and adding EG00229 and VEGF164 together increased extravasation more than adding VEGF164 alone (**Fig. 1A,B**). These findings show that EG00229, albeit able to inhibit VEGF164-dependent vascular permeability, can also activate it independently of VEGF164. Therefore, we characterised the EG00229 response in more detail. Re-treatment with VEGF164 and EG00229 after 15 min washout induced dye extravasation again without altering efficacy, even when VEGF164 and EG00229 treatment was alternated (**Fig. S1C**). These findings suggest that both VEGF164 and EG00229 can increase retinal vascular permeability transiently and reversibly. Although several studies have used EG00229 at 10 µM [19–21], others have used 30 µM [22], 100 µM [18,13] or 2 µM [20]. We have routinely used 10 µM EG00229, but EG00229 also induced retinal vascular leakage at 10-fold lower or 10-fold higher concentrations (**Fig. 1C**). As leakage appeared maximal at 10 µM (**Fig. 1C**), this dose was used in subsequent experiments. Together, these findings suggest that EG00229 is a CendR mimetic that prevents VEGF-induced vascular permeability in retinal microvasculature, but also increased it on its own, similar to VEGF164.

**Figure 1.**
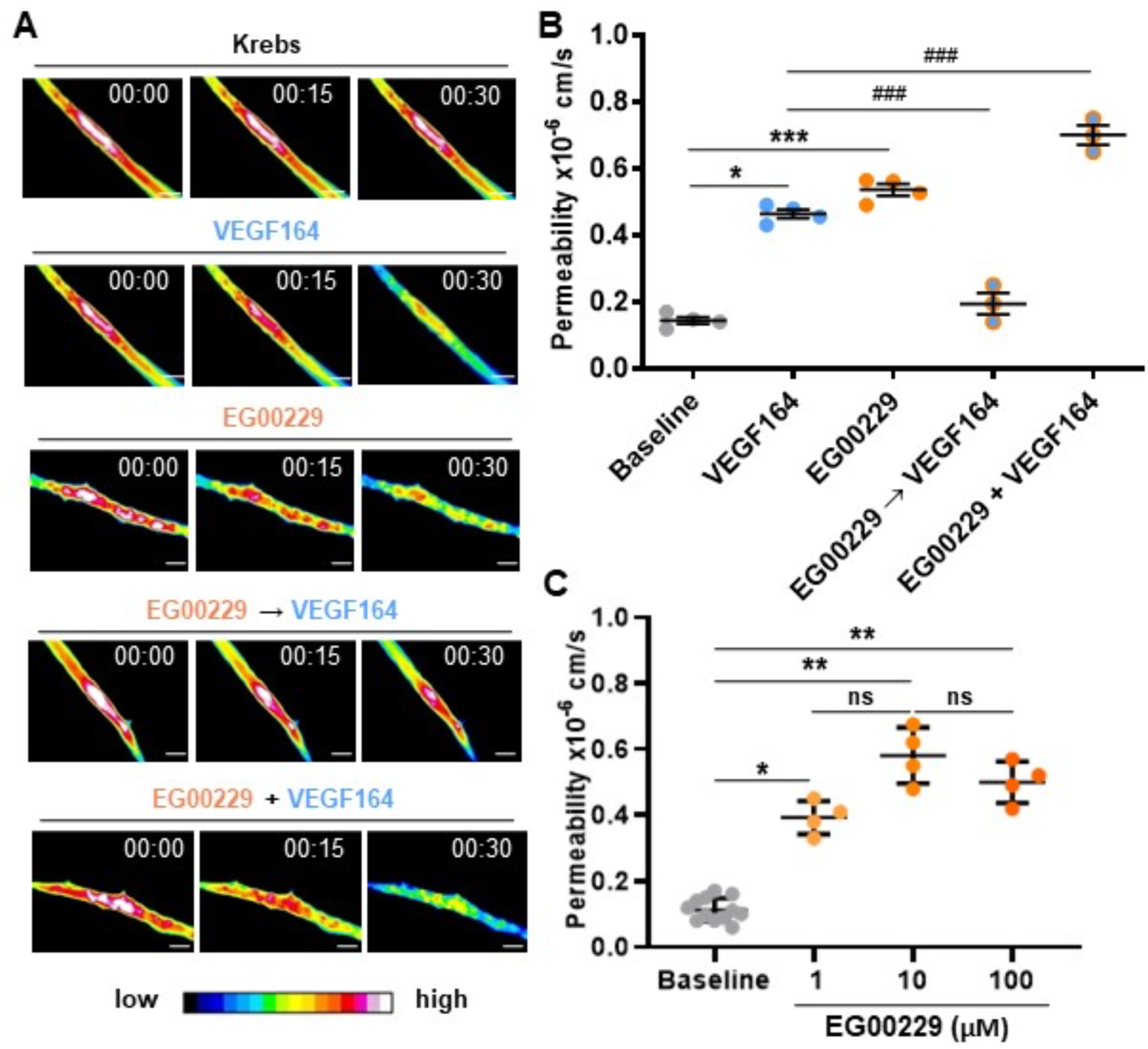
EG00229 induces retinal vascular leakage. (**A**) Representative 16-color pseudocoloured images of fluorescence intensity in vessel segments of wild type *ex vivo* retina at the indicated times after treatment with vehicle (Krebs) or VEGF164 and/or EG00229 as indicated. (**B,C**) Quantification of fluorescence changes over 2 min compared to baseline (vehicle), after (**B**) treatment with EG00229 (n = 4), VEGF164 (n = 4), VEGF164 after EG00229 pre-treatment (n = 3) or co-treatment of EG00229 and VEGF164 (n = 3), or after (**C**) treatment with different EG00229 concentrations (n = 4 each). Data are shown as mean ± SD. Each data point indicates the value for one retina from one mouse after one instance of adding a test substance. Repeated measures one-way ANOVA followed by post-hoc Dunnett’s test; asterisks: significant P-values for test substance compared to baseline: * P < 0.05, ** P < 0.01,*** P < 0.001; hashtags: significant P-values for test substance(s) compared to VEGF164: ### P < 0.001; ns, not significant (P > 0.05).

### EG00229 induced retinal vascular leakage through NRP1 but not VEGFR2 or VEGFR1

Prior studies showed that NRP1 promotes VEGF164-induced vascular permeability in the dermis together with VEGFR2 [30]. We, therefore, examined whether EG00229 also induced vascular leakage in the retina of mice after endothelial knockdown of VEGFR2, which is encoded by *Kdr*. This, we administered tamoxifen to adult *Kdr^fl/fl^*mice expressing or lacking *Cdh5*-CreER^T2^ [30]. In these experiments, knockdown in adulthood, after completion of vascular development, ensured normal retinal vascular density. Immunoblotting confirmed VEGFR2 knockdown (**Fig. 2A**). Agreeing with pharmacological VEGFR2 inhibition [31], VEGF164-induced dye extravasation in retinal explants was abolished by VEGFR2 knockdown, whereas EG00229-induced dye extravasation was not (**Fig. 2B**). As VEGFR1 interacts with NRP1 in vitro [38] and promotes VEGF signalling together with NRP1 during psoriasis-like disease [39] and in snake venom-induced dermal vascular leakage [40], we next examined how endothelial VEGFR1 knock-down affected retinal vascular leakage by targeting the VEGFR1 gene, *Flt1*. Thus, we administered tamoxifen to adult *Flt1^fl/fl^*mice expressing or lacking *Cdh5*-CreER^T2^ and confirmed VEGFR1 knockdown (**Fig. 2C**), but found that it did not impair VEGF164- or EG00229-induced dye extravasation (**Fig. 2D**). By contrast, endothelial NRP1 knockdown significantly reduced both VEGF164- or EG00229-induced dye extravasation (**Fig. 2E,F**). Notably, NRP1 knockdown did not abrogate VEGF164-induced leakage completely (**Fig. 2F**), which agrees with prior findings in the dermis, where VEGFR2 is absolutely required but NRP1 enables a robust response [30]. Compared to VEGF164, dye extravasation induced by EG00229 was affected more severely by endothelial NRP1 knockdown (**Fig. 2F**; the reduction in extravasation in individual mice correlated with knockdown efficiency). This finding shows that NRP1 is required for EG00229-induced retinal vascular leakage, thereby demonstrating that this response is not an off-target effect.

**Figure 2.**
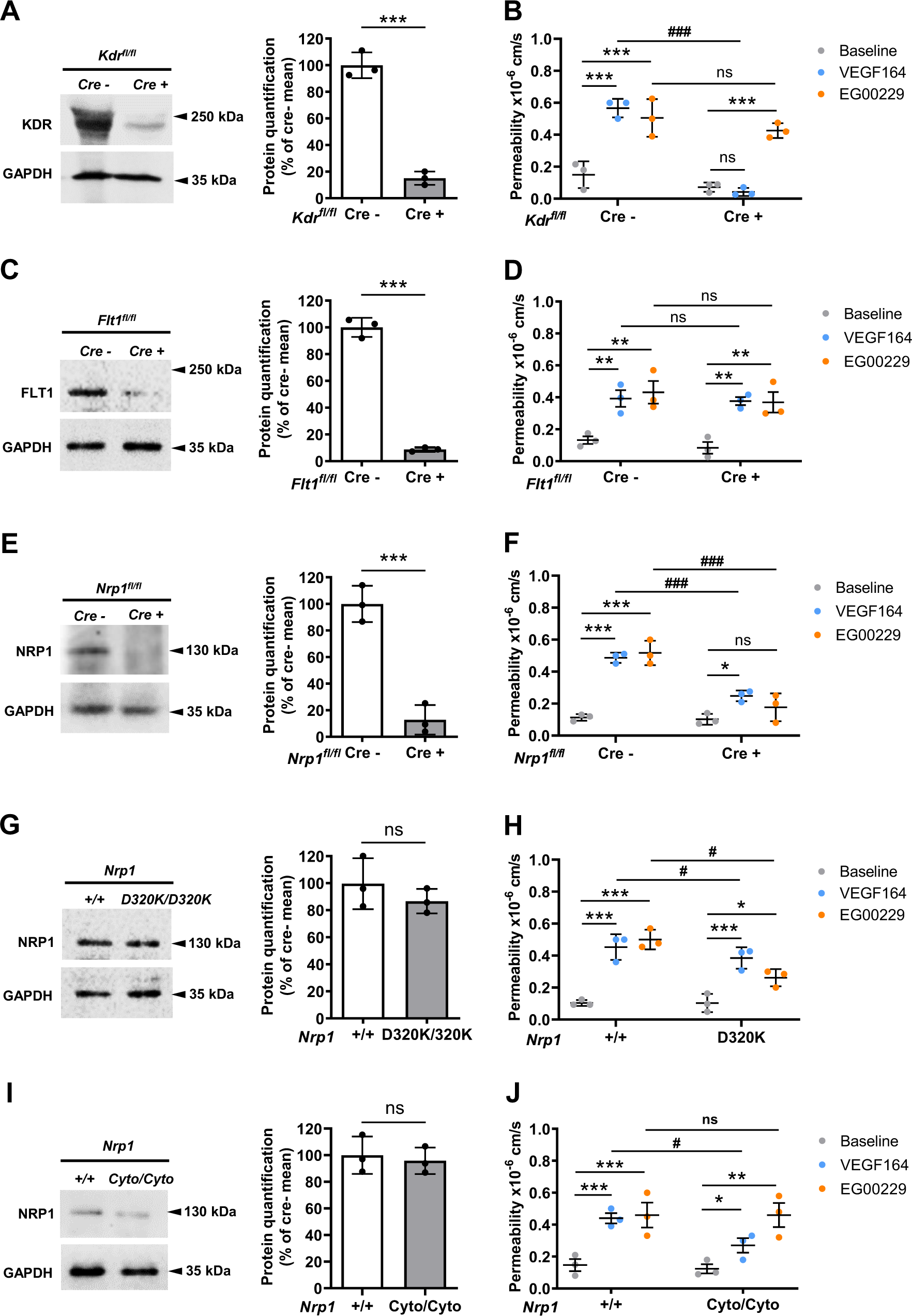
EG00229 induces retinal vascular leakage via NRP1, independently of KDR and FLT1. (**A-F**) EG00229-induced retinal vascular leakage depends on endothelial NRP1, but not VEGFR2 or VEGFR1. Adult *Kdr^fl/fl^* (**A,B**), *Flt1^fl/fl^* (**C,D**) and *Nrp1^fl/fl^* (**E,F**) mice expressing or lacking *Cdh5*-CreER^T2^ were tamoxifen-treated and used two weeks after the last tamoxifen dose for lung immunoblotting with the indicated antibodies (**A,C,E**) and *ex vivo* retina permeability assays (**B,D,F**). (**G-J**) EG00229-induced retinal vascular leakage depends on NRP1’s CendR but not cytoplasmic domain. Immunoblotting of lung lysates with the indicated antibodies (**G,H**) and *ex vivo* retina permeability assays (**I,J**) in *Nrp1^D320K/D320K^* (**E,F,H**) and *Nrp1^cyto/cyto^*(**I,J**) mice. In all experiments, mice were used as tissue donors for parallel lung immunoblotting to determine receptor protein levels and permeability assays with one retina per mouse to determine the genotype effect on VEGF164-versus EG00229-induced vascular leakage; n = 3 mice for each genotype (the second retina was used for experiments shown in Figure 3). The migration of the molecular weight marker is indicated with arrowheads to the left of each immunoblot. Lung immunoblotting: Data are shown as mean ± SD; each data point indicates the value for one retina from one mouse. Asterisks indicate significant P-values for changes in the level of the indicated proteins (unpaired Student’s t-test): *, P < 0.05; **, P < 0.01; ***, P < 0.001; ns, not significant (P > 0.05). *Ex vivo* retinal permeability assay: For repeated testing of the same retina, the order of adding VEGF164 or EG00229 was alternated, with washing each time in Krebs solution for 10 min followed by recording the new baseline. Data are shown as mean ± SD; each data point indicates the value for one retina from one mouse, whereby all recordings for one agent in one retina were averaged to obtain the value for that retina. Significant P-values are shown for fluorescence changes induced by VEGF164 or EG00229 versus Krebs for specific genotypes (asterisks) and in mutants versus controls (hashtags: *, P < 0.05; **, P < 0.01; ***, P < 0.001; #, P < 0.05; ##, P < 0.01; ns, not significant (P > 0.05). Repeated measures two-way ANOVA.

### EG00229 induced retinal vascular leakage via NRP1’s CendR pocket but not the NCD

Next, we analysed *Nrp1^D320K/D320K^* mice in which the D320 residue in NRP1’s CendR pocket is replaced with K [30,41]. These mice have normal NRP1 levels (**Fig. 2G**), but impaired VEGF164 binding to NRP1 [41]. In analogy to findings in the dermis [30], VEGF164-induced retinal vascular leakage was attenuated in *Nrp1^D320K/D320K^* mice (**Fig. 2H**). EG00229-induced retinal vascular leakage was also significantly reduced in these mice (**Fig. 2H**), again corroborating that this response is not an off-target effect but a NRP1-dependent, CendR pocket-mediated mechanism.

Prior work showed that NRP1’s NCD synergises with NRP1’s VEGF binding domain to promote complex formation between VEGFR2 and NRP1 to enhance VEGF164-induced VEGFR2 signalling [42,43]. In agreement, *Nrp1^cyto/cyto^* mice lacking the NCD have reduced VEGF164-induced vascular leakage in the dermis [30]. We confirmed that NRP1 protein in *Nrp1^cyto/cyto^* mice was stable despite lacking the NCD (**Fig. 2I**). Although VEGF164-induced dye extravasation was attenuated in *Nrp1^cyto/cyto^* mice, EG00229-induced dye extravasation was not (**Fig. 2J**). The finding that EG00229 did not require the NCD to induce vascular leakage thus distinguished its mechanism of action from that of VEGF164 and a multivalent RPARPAR peptide, which both require the NCD.

### VEGF165 and EG00229 bind similarly to NRP1’s b1 domain

To better understand why EG00229 has the capacity to signal via NRP1, similar to VEGF164, we compared the published X-ray crystal structures of NRP1’s b1 domain in complex with VEGF164 versus EG00229 [18,44,41,33]. Thus, we used PyMol and LigPLOT+ to compare VEGF164’s exon 7-8 domain (PDB:4deq) versus EG00229 (PDB:3i97), which illustrated that both ligands interact in a similar manner with NRP1’s CendR pocket (**Fig. 3A,B**). Firstly, they form salt bridges with D320 via VEGF165’s terminal arginine EG00229’s arginyl moiety (**Fig. 3C,D**) and hydrogen bonds with S346, T349 and Y353. VEGF165’s exon 7 has additional interactions with NRP1’s L1 loop via T299 and N300 (**Fig. 3C**), whose replacement with analogous residues from NRP2 reduces VEGF165 binding [33]. Secondly, EG00229 interacts with NRP1’s L1 loop, similar to VEGF165, via a stacking interaction between the benzothiadiazole ring and N300 [18] and forms a hydrogen bond with S298 that has not previously been commented on (**Fig. 3D**). Y297 appears to contribute to shaping the binding pocket via nonpolar interactions with VEGF165 and EG00229 and an additional hydrogen bond with VEGF165 that is absent with EG00229 (**Fig. 3C**). The striking similarity in NRP1’s b1 interactions with VEGF165 and EG00229 raised the possibility that EG00229 binding to the CendR pocket might activate similar NRP1-dependent signalling pathways as those known to mediate VEGF164-induced vascular permeability.

**Figure 3.**
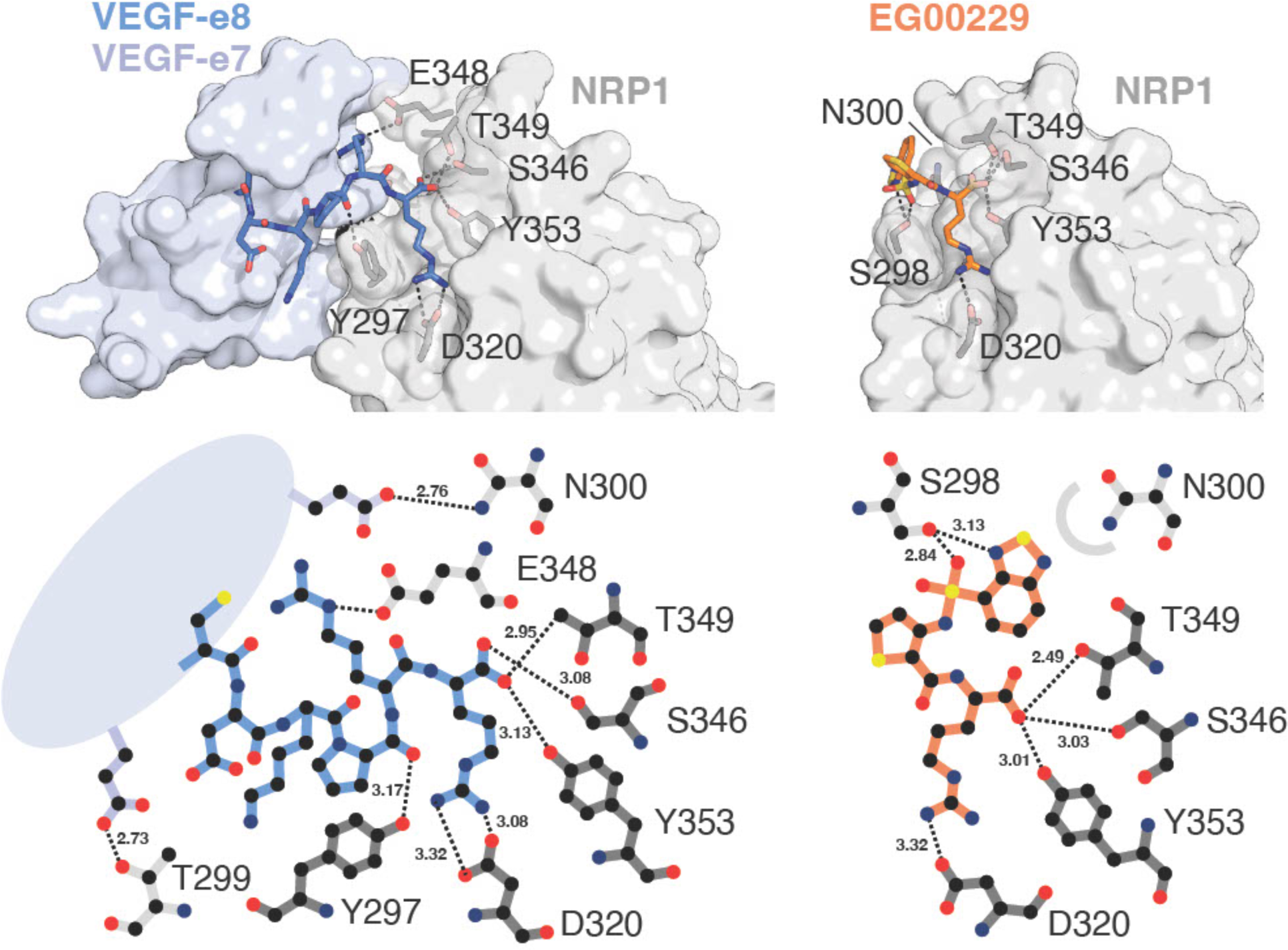
VEGF165 and EG00229 bind similarly to the NRP1 b1 domain. Schematic representation of the crystal structures (**A,B**) and corresponding interaction diagrams (**C,D**) of the NRP1 b1 domain in complex with VEGF165 exon 7-8 (PDB:4deq; **A,C**) or the NRP1 b1 domain with EG00229 (PDB:3i97; **B,D**). Key interacting NRP1 residues are shown with stick representation with corresponding hydrogen bonds as black dashes. A stacking interaction is represented by a semi-circle in (**B)**.

### EG00229 induces retinal vascular permeability via SFK-induced p38 activation

Next, we asked whether EG00229 induced endothelial signalling pathways implicated in retinal vascular permeability. To begin, we examined T180/Y182 phosphorylation of p38 (encoded by *Mapk14*), which is required for VEGF164-induced vascular leakage in the retina and brain [31,32]. Retinal wholemount immunostaining showed that EG00229 induced endothelial p38 T180/Y182 phosphorylation (P-p38), similar to VEGF164 (**Fig. 4A**). VEGFR2 knockdown attenuated VEGF164-induced endothelial P-p38, but not EG00229-induced endothelial P-p38 in retinal vasculature (**Fig. 4A**), and VEGFR1 knockdown neither affected EG00229-nor VEGF164-induced endothelial P-p38 in retinal vasculature (**Fig. 4B**). NRP1 knockdown and the D320K mutation attenuated both VEGF164- and EG00229-induced P-p38 in retinal vasculature (**Fig. 4C,D**). These results demonstrate that EG00229 depends on NRP1 but not VEGFR1 or VEGFR2 for inducing a key vascular permeability signalling pathway in the retina, and they corroborate our findings in the retinal vascular leakage assay to show that the receptor requirement for VEGF164 and EG00229 differs with respect to VEGFR2.

**Figure 4.**
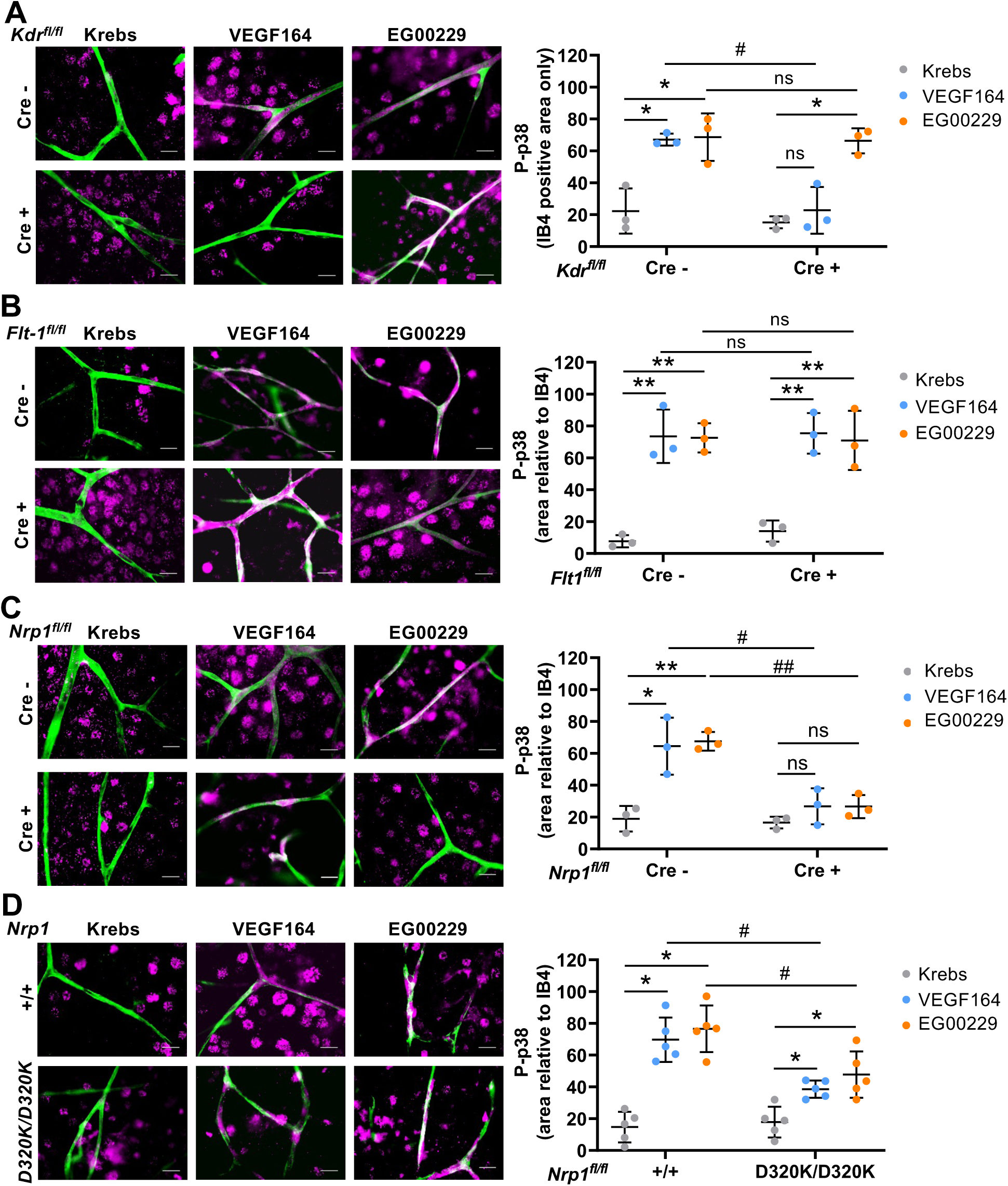
EG00229 activates p38 in retinal vasculature through NRP1’s, independently of KDR and FLT1. Freshly dissected retinae from mice of the indicated genotypes were incubated in Krebs solution with or without VEGF164 or EG00229, fixed and immunostained with the vascular EC marker isolectin B4 (green) and an antibody against P-p38 (magenta). tinas were obtained from the second eye of mice used for *ex vivo* permeability experiments in Figure 2. In (**A-C**), we used retinas from adult *Kdr^fl/fl^* (**A**), *Flt1^fl/fl^* (**B**) and *Nrp1^fl/fl^* (**C**) mice expressing or lacking the endothelial *Cdh5*-CreER^T2^ transgene, which had been tamoxifen-treated two weeks before *ex vivo* treatment of the retina. In (**D**), we used retinas from *Nrp1^D320K/D320K^* and wild type littermate mice for *ex vivo* treatment. Re Epifluorescent images (scale bars: 10 μm) show P-p38 in IB4-positive vasculature and IB4-negative neuroglial cells. Epifluorescent images were then used for quantification of P-p38 pixel intensity in the IB4-positive area. Data are shown as mean ± SD; each data point represents the value for one retina from one mouse; n = 3 mice for each genotype. Significant P-values are shown for P-p38 staining area for VEGF164 or EG00229 versus vehicle (Krebs) for each genotype (asterisks) and for VEGF164 or EG00229 in mutant versus control (hashtags) (two-way ANOVA): *, P < 0.05; **, P < 0.01; ***, P < 0.001; #, P < 0.05; ns, not significant (P > 0.05).

To further probe the signalling requirements for EG00229-induced endothelial signalling, we used established small molecule inhibitors. First, we used SU1498 as an established inhibitor of VEGF164-induced retinal vascular leakage [31] to determine whether a pharmacological approach recapitulated our findings with EG00229 in genetic knockout mice. Agreeing with prior work, pre-incubating the retina with SU1498 abolished sulforhodamine B extravasation induced by VEGF164 (**Fig. 5A**). By contrast, pre-incubating the retina with SU1498 did not diminish EG00229-induced dye extravasation (**Fig. 5A**). Therefore, a pharmacological approach corroborated that EG00229 does not require VEGFR2 to induce vascular leakage. Second, we used the p38 inhibitor SB203580, which decreases VEGF164-induced retinal vascular leakage [31]. SB203580 also significantly decreased EG00229-induced dye extravasation (**Fig. 5B**). Third, we used PP2 [46], which in dermal ECs disrupts VEGF164-induced permeability signalling [30]. Treatment with PP2 at 10 µM, which inhibits both SFK and p38 activation {Bain, 2003 #6076}, also effectively abolished EG00229-induced dye extravasation in the retina (**Fig. 5C**). Further, wholemount retinal immunostaining showed that EG00229-induced endothelial p38 activation was not impaired by SU1498 but SB203580 and PP2 (**Fig. 5D**).

**Figure 5.**
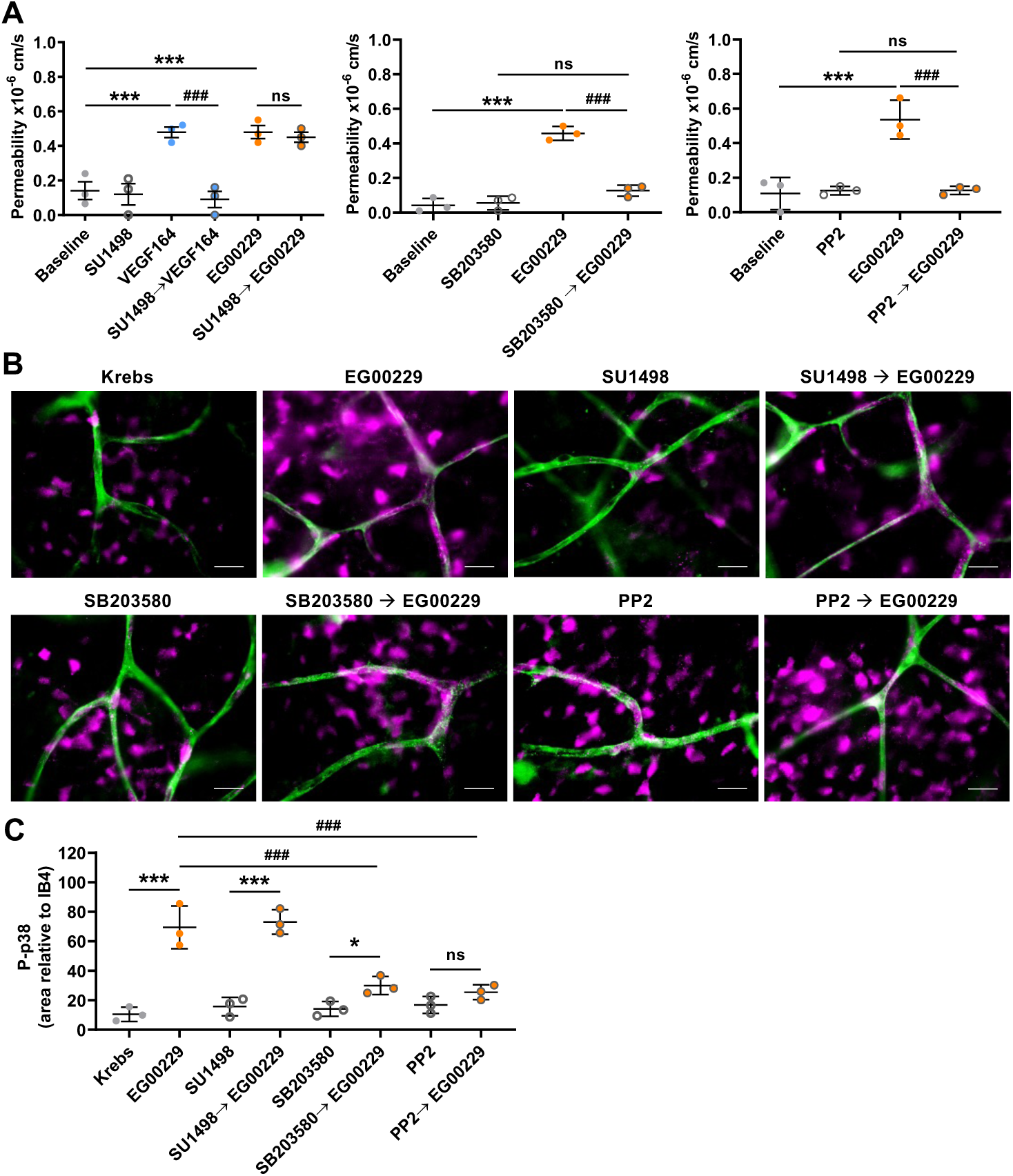
EG00229 induces retinal vascular leakage via p38 and SFK activation. (**A**) Quantification of sulforhodamine B fluorescence changes in the *ex vivo* retina permeability assay after treatment with SU1498, SB203580 or PP2 (inhibitors) alone, compared to VEGF164 or EG00229 treatment with or without inhibitor pre-treatment (n = 3 treatments of 3 independent retinas each). Data are shown as mean ± SD; each data point represents the value for one retina from one mouse. Significant P-values are shown for inhibitor versus baseline (asterisks) and EG00229 ± inhibitor (hashtags): *** P, < 0.001; ###, P < 0.001; ns, not significant (P > 0.05). Repeated measures one-way ANOVA. (**B,C**) Freshly dissected retinae from wild type mice were incubated in Krebs solution (vehicle) with or without EG00229, or preincubated with SB203580 or PP2 for 15 min before adding EG00229. *Ex vivo* retinae were then fixed and immunostained with the vascular EC marker isolectin B4 (IB4, green) and an antibody against P-p38 (magenta). Epifluorescent images (**B**, scale bars: 10 μm) were used for quantification of pixel intensity for P-p38 in the IB4-positive vascular area (**C**). Significant P-values for P-p38 staining in the IB4+ area for inhibitors (SB203580 or PP2 versus SB203580) versus EG00229 are indicated with asterisks or after pre-treatment with the inhibitors, indicated with hashtags: *, P < 0.05; ***, P < 0.001; ###, P < 0.001; ns, not significant (P > 0.05); one-way ANOVA.

Together, these findings demonstrate that p38 conveys EG00229-induced signals for retinal vascular leakage.

### EG00229 induce paracellular permeability signalling in brain microvascular ECs

We next asked whether EG00229 also increased the permeability of brain vascular endothelium. For this experiment, we grew primary microvascular ECs isolated from rat brains on transwells without passaging, because this method preserves excellent junctional barrier properties for flux studies as a proxy of extravasation from microvessels [32].Thus, we measured fluorescent dextran flux across these brain endothelial monolayers (**Fig. S2A**). As shown [32,31], VEGF164 increased flux across the monolayer by nearly 2-fold (**Fig. S2B,C**). Similarly, EG00229 enhanced flux across the monolayer by nearly 2-fold (**Fig. S2B,C**).

To increase vascular leakage in brain ECs, VEGF164 activates p38 and SFK signalling [31,30,47]. Immunoblotting showed that EG00229 activates p38 in brain endothelial monolayers, which peaked around 5 min and returned to baseline 30 min after treatment (**Fig. 6A,B**). Further, immunoblotting with an antibody that detects tyrosine phosphorylation of the SFK catalytic domain (corresponding to Y419 in human SRC) showed that EG00229 also activates SFKs in brain endothelial monolayers with similar kinetics (**Fig. 6A**), and pre-incubating rat brain EC monolayers with PP2 significantly decreased EG000229-induced P-p38 (**Fig. 6C**). To investigate relevance for human brain EC, we used hCMEC/D3 cells, which are used to model the human blood-brain barrier [48,37]. Again, we found that EG00229 induced both p38 and SFK activation in these cells (**Fig. S2D**). The ability of EG00229 to induce vascular leakage signalling is therefore conserved from rodent to human brain ECs.

**Figure 6.**
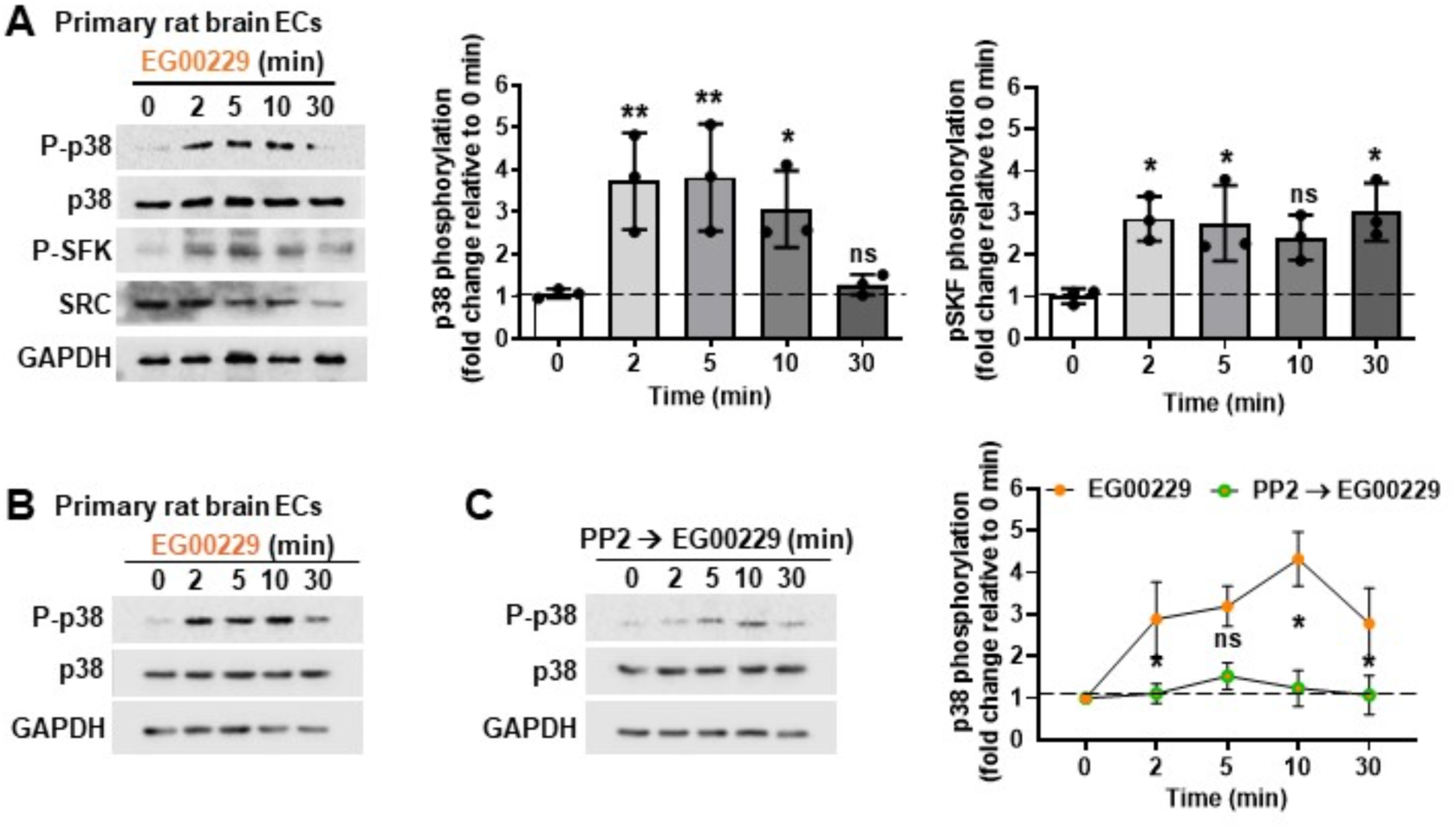
EG00229 induces paracellular permeability signalling in rat brain ECs. Confluent primary rat brain ECs were treated with EG00229 for the indicated times. Cell lysates were used for immunoblotting with the indicated antibodies, followed by quantification of pixel intensities for P-p38 and P-SFK relative to total p38 and SRC pixel intensities, respectively. GAPDH was used as a loading control. In (**A**), p38 and SFK phosphorylation levels after EG0229 treatment were quantified at different time points relative to 0 min. In (**B**), EG00229 treatment was preceded by incubation with PP2; decreased EG00229-induced p38 phosphorylation after PP2 treatment was compared to EG00229 alone at each time point. Data are shown as mean fold change ± SD. Asterisks indicate significant P-values for phosphorylation induction after EG00229 treatment (: *, P < 0.05; **, P < 0.01; ns, not significant (P > 0.05); one-way ANOVA.

To increase vascular leakage, VEGF164 acts on intercellular junctions in a mechanism termed paracellular permeability, including in brain ECs [32]. This pathway involves VEGFR2-induced signal transduction events that likely modify the adherens junction component vascular endothelial (VE) cadherin (CDH5) by phosphorylation [49] and nitrosylation [50]; these modifications are reported to cause CDH5 redistribution in HUVEC cultures from a linear to a ‘zig-zag’ pattern [51], and in brain EC cultures from a linear pattern to focal broadening especially at tricellular junctions [31]. As previously shown [31,52], CDH5 localised in such a broader pattern around contact sites in VEGF164-treated compared to untreated rat brain EC monolayers (**Fig. 7A**). Similar overall CDH5 staining intensity before and after VEGF164 treatment agrees with CDH5 not being degraded after VEGF164 treatment [49,50], but that modifications caused CDH5 redistribution [51]. A similar CDH5 redistribution, also without changes in staining intensity, occurred after EG00229 treatment of monolayers (**Fig. 7B**). The broader CDH5 localisation after EG00229 treatment was particularly obvious at tricellular junctions (**Fig. 7B**), as previously reported for VEGF164 treatment of brain EC monolayers [31]. Together with the observed changes in SFK and p38 activation, CDH5 re-distribution in brain ECs suggests that EG00229 increases paracellular permeability via pathways that are similar to those induced by VEGF164, despite not needing VEGFR2 activation (**Fig. S3**).

**Figure 7.**
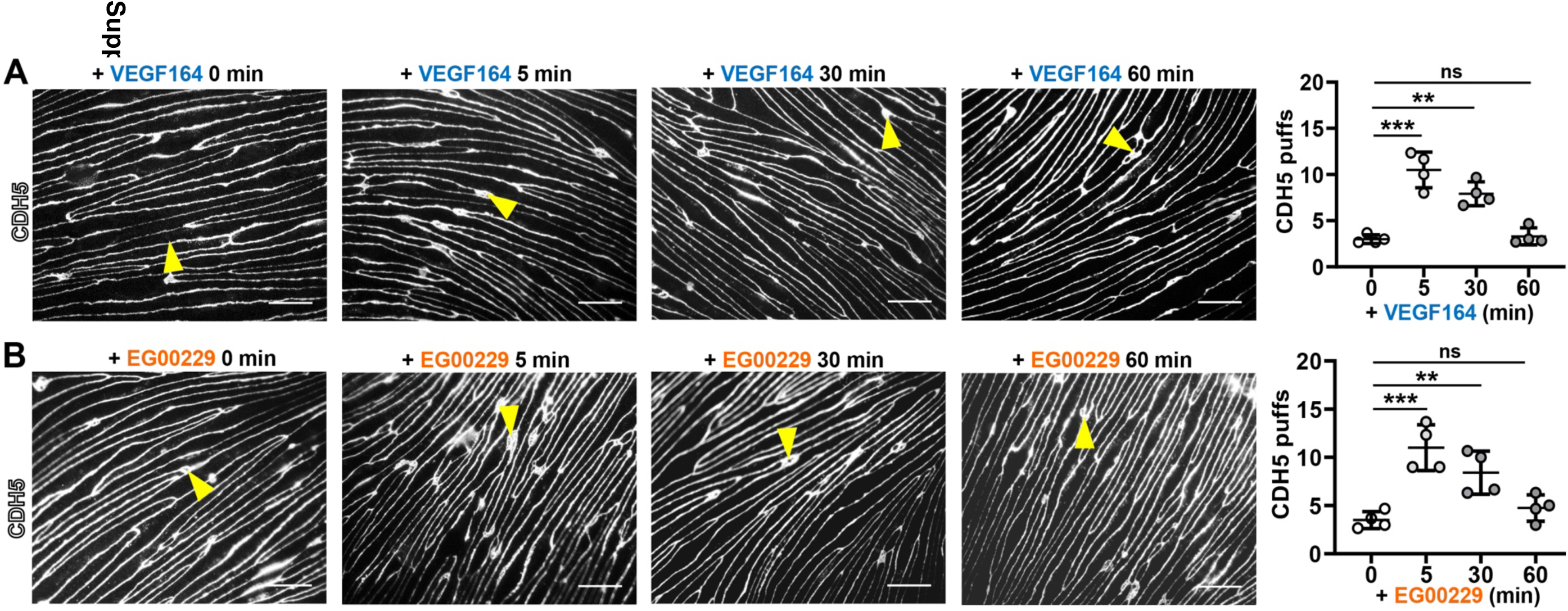
EG00229 causes junctional remodelling in brain microvascular EC monolayers. Epifluorescence images of confluent primary rat brain ECs for CDH5 fixed were immunostained at the indicated times after VEGF164 (**A**) or EG00229 (**B**) treatment; scale bars: 100 μm. Yellow arrowheads indicate broad CDH5 staining (‘puffs’). Quantification of puff number: data show the mean ± SD from 4 independent primary cell preparations; each data point represents the average of 3 transwells (technical repeats). Asterisks indicate significant P-values for the number of puffs compared to 0 min (one-way ANOVA): ** P < 0.01; *** P < 0.001; ns, not significant (P > 0.05

## Discussion

EG00229 was originally designed to inhibit VEGF164-induced NRP1 responses [18]. Accordingly, EG00229 pre-treatment inhibits VEGF164-binding to NRP1 with an IC_50_ of 8 µM in vitro [18]. As NRP1 promotes vascular hyperpermeability in mouse models of ocular neovascular disease [30,24], we asked whether EG00229 was effective in inhibiting VEGF164-induced retinal vascular leakage. Indeed, 10 µM EG00229 attenuated VEGF164-induced sulforhodamine B leakage in the *ex vivo* mouse retina model of vascular permeability (**Fig. 1**). However, adding EG00229 alone as a control in our experiments also induced acute vascular leakage it (**Fig. 1**). Whereas EG00229’s ability to inhibit VEGF164 binding to NRP1 has been studied extensively, we are, to the best of our knowledge, the first to identify that EG00229 can actively induce a biological response. Our findings, therefore, suggest that testing EG00229 in the absence of another CendR ligand is an important control when establishing its ability to inhibit NRP1-mediated responses. More broadly, our findings suggest that drugs designed to bind NRP1’s CendR pocket should be routinely screened for potential effects on vascular permeability at neurovascular barriers.

Prior work showed that a multimerised CendR peptide (RPARPAR) and a NRP1 extracellular domain-binding antibody induce vascular leakage via NRP1, independently of VEGFR2 [12,11]. To explain VEGFR2 independence, it has been hypothesised that CendR peptide-induced NRP1 clustering stimulates NRP1 internalisation, because internalisation or vascular leakage were more effectively induced by multivalent than monomeric RPARPAR peptide [11]. The small molecule EG00229 lacks obvious molecular features present in multivalent agents that cluster NRP1, and, therefore, it remains elusive how EG00229 binding to NRP1’s CendR pocket induces vascular leakage. To better understand VEGF165 versus EG00229 binding to NRP1, we directly compared prior knowledge of their structural interactions with NRP1 (**Fig. 3**). Notably, the published X-ray crystallography structures of VEGF165 versus EG00229 bound to NRP1 [18,33,41] showed similar interactions between the ligands and NRP1’s b1 domain (**Fig. 3**). Thus, the D320 requirement for NRP1 binding has been demonstrated for VEGF165 [41,30] and, here, also for EG00229 (**Fig. 2**). Also similar to VEGF1645, EG00229 interacts with N300 in NRP1’s L1 loop [18,33] (**Fig. 3**). Similar interactions with the L1 loop are not seen with other CendR-type ligands, such as tuftsin [53] and the KDKPPR peptide [54]. Conceivably, therefore, EG00229 might differ from other CendR-like NRP1 targeting compounds by being particularly adept at mimicking the VEGF165 C-terminus. EG00229’s dependence on NRP1 for leakage induction was confirmed in endothelial NRP1 knockout mice and in *Nrp1^D320K/D320K^* mice lacking a fully functional CendR pocket (**Fig. 2**). Whereas endothelial NRP1 loss completely abolished leakage induction by EG00229, the D320K mutation did not abolish it completely. Therefore, NRP1 is required to convey EG00229’s effect, and the D320 residue is important for their interaction, confirming the specificity of EG00229’s activity for NRP1. Further mutation of other interacting residues may be required to completely disrupt the docking of CendR ligands to NRP1’s binding pocket.

To better define the molecular pathways through which EG00229 induced acute retinal vascular leakage, we established EG00229’s signalling requirements. Consistent with EG00229 lacking a VEGFR2 binding site, genetic or pharmacological VEGFR2 ablation did not affect EG00229-induced sulforhodamine B extravasation from retinal vessels (**Fig. 2**). This finding is reminiscent of those made for the aforementioned RPARPAR CendR peptide and a NRP1 extracellular domain-binding antibody that did not require VEGFR2 to induce leakage of dermal vessels [12,11]. VEGFR2 independence of EG00229-induced retinal vascular leakage and p38 activation was not explained by a VEGFR1 role (**Figs. 2, 4**), even though VEGFR1 promotes dermal vascular leakage in response to VEGF164 [40]. Taken together with prior results for the RPARPAR peptide, it appears that VEGF receptor tyrosine receptor activation is not a general feature of NRP1’s CendR ligands but is important for VEGF164, which has both a CendR peptide domain and a VEGFR binding domain.

We further show that the mechanisms of EG00229-induced retinal vascular leakage was independent of NRP1’s highly conserved intracellular domain, the NCD (**Fig. 2**) and therefore differed from that of the multimerised CendR peptide RPARPAR and VEGF164, which both require the NCD to induce vascular leakage [12][30]. In addition to providing knowledge of EG00229 signalling, our findings also contribute new knowledge of VEGF signalling. Thus, the receptor requirement for VEGF164-induced vascular permeability responses in retinal permeability assays is similar to that in dermal permeability assays with respect to VEGFR2 and NRP1, but different with respect to VEGFR1, which contributes to VEGF164-induced dermal but not retinal vascular leakage. Finding that VEGFR1 was not required for VEGF164-induced retinal vascular leakage agrees with prior findings that VEGFR1 localises preferentially to the luminal side of brain and retina endothelium (*35*), rather than the tissue-facing side of the endothelium that is exposed to VEGF both in vivo and in our ex vivo assays.

To investigate whether EG00229 differs from RPARPAR or VEGF164 by differentially activating downstream signal transducers, we have examined P38 and SFK phosphorylation. This was important, because multimerised RPARPAR reportedly does not induce VEGFR2 or P38 activation and therefore differs from VEGF164 [12]. EG00229 activated p38, and inhibiting p38 activation with SB203580 attenuated EG00229-induced vascular leakage (**Fig. 5**). EG00229 therefore differed from multimerised RPARPAR [12], but acted similar to VEGF164 with respect to NRP1-dependent p38 activation [57,32,31,58], despite lacking obvious means of engaging VEGFR1 or VEGFR2 to activate downstream phosphorylation cascades. Next, we considered that VEGF-induced paracellular permeability signalling depends on VEGF164-induced SFK activation [59,45,60,47], which is NRP1-dependent [30]. Similar to VEGF164, EG00229 induced SFK activation (**Fig. 6** and **S2**) suggests that assaying linked SFK and p38 phosphorylation might be useful when screening NRP1-targeting compounds for their ability to modulate the neurovascular barrier, particularly when evaluating CendR-based drugs intended to treat human disease. Further, observing similar permeability signalling in mouse, rat and human ECs in response to EG00229 (**Figs. 4, 6** and **S2**) corroborates that rodent ECs are likely relevant for modelling signalling events at the neurovascular barrier, as proposed [37,48,61].

Although chronic increases in vascular leakage pose a significant clinical problem in ischemic conditions of the brain and retina, the *ex vivo* retina model measures specifically acute paracellular permeability induction. Further work is thus needed to determine whether cumulative effects of prolonged EG00229 treatment would translate into clinically relevant neurovascular barrier modulation, such as oedema due to chronic VEGF164 upregulation. Thus, it should be investigated whether EG00229’s ability to open the neurovascular barrier might enhance drug penetration into cells or across vascular barriers, in analogy to the benefit of multivalent CendR peptide-coated particles, which promote the transportation of payloads deep into extravascular tissue [11].

## Statements and Declarations

The authors have no competing interests to declare.

## Contributions

SD, AF, JTB, PT and CR conceived and designed the study. SD performed most of the experiments. JTB, AF and PT performed some of the experiments. CB prepared the schematic representation of the crystal structures and corresponding interaction diagrams. SD and AF analysed data. LD bred and genotyped mice. SD and CR co-wrote the manuscript. All authors have read, commented on and approved the submitted manuscript.

## Acknowledgements

We thank the Biological Resources team and Yaiza Yuste Herranz for technical help. This study was supported by research grants from the Wellcome Trust (095623/Z/11/Z), British Heart Foundation (PG/14/81/31119, PG/17/70/33232) and MRC (MR/N011511/1) to CR, British Heart Foundation (FS/13/59/30649) to JTB, Diabetes UK (Ref 17/0005637) to PT and SD and a Moorfields Eye Charity Career Development Award (GR001458) to SD.

---

**Figure S1.**
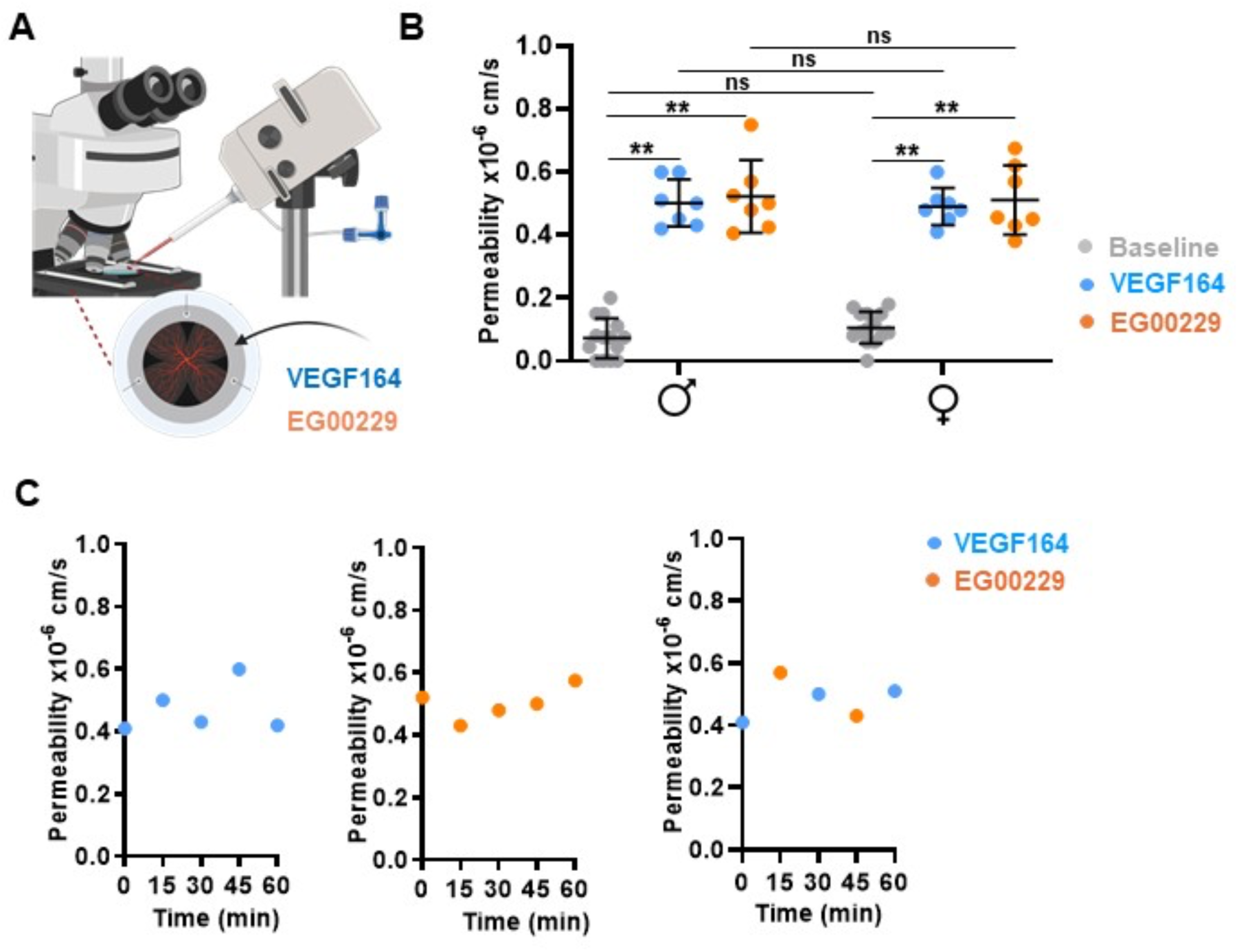
EG00229 increases retinal microvascular EC permeability similar to VEGF164. (**A**) Schematic representation of the *ex vivo* retinal permeability assay. (**B,C**) Quantification of retinal vascular leakage in the *ex vivo* retinal vascular permeability assay after adding EG00229 or VEGF164 in retinae of (**B**) male and female wildtype mice (n = 7 each) and (**C**) after repeated addition of VEGF164 and EG00229 to the same retinal vessel, or when added alternately; for repeated testing, the retina was washed with Krebs for 10 min and a new baseline recorded. Data are shown as mean ± SD. Each data point indicates the value for the fluorescence change in one retina from one mouse obtained 2 min after one instance of adding a test substance. Repeated measures two-way ANOVA followed by post-hoc Dunnett’s test: asterisks indicate significant P-values for test substance addition compared to baseline (Krebs); *, P < 0.05; **, P < 0.01; ***, P < 0.001; ns, not significant (P > 0.05).

**Figure S2.**
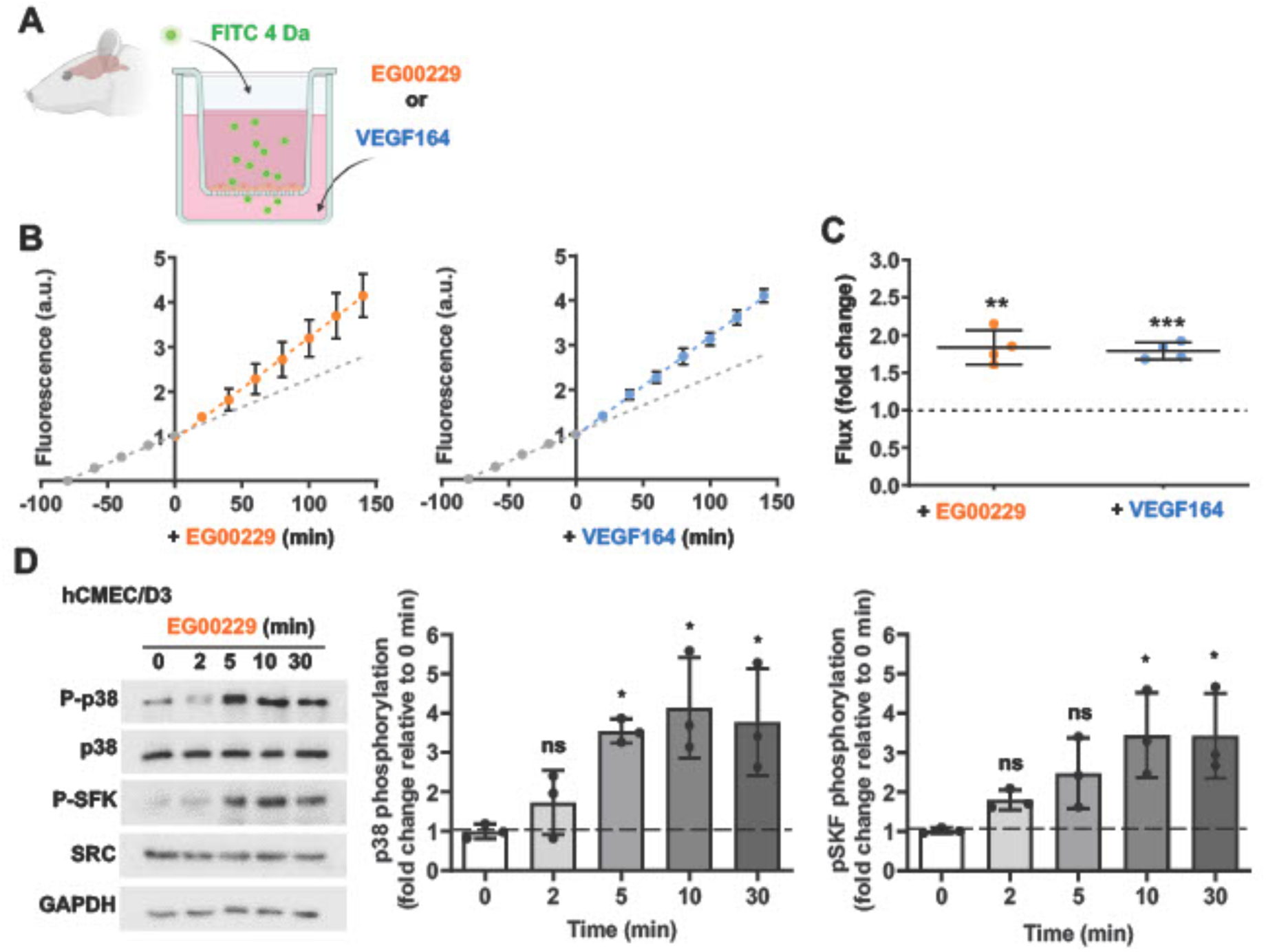
EG00229 increases brain microvascular EC permeability and activates p38 and SKF. (**A**) Schematic of the transendothelial flux assay with brain microvascular ECs. Primary rat brain microvascular ECs were grown on permeable transwell inserts until they reached a TEER greater than 200 Ω/cm2. 4 kDa FITC dextran was then added to the apical chamber. Baseline FITC flux was recorded as the time-dependent accumulation of fluorescence in the basal chamber before adding EG00229 or VEGF164 and continued recording. (**B,C**) EG00229 increases the permeability of brain microvascular EC monolayers. Linear regression analysis of flux over time before and after adding VEGF164 or EG00229, defined as time 0. (**B**) Data are shown as mean ± SD flux change from 4 independent experiments over time. (**C**) Data are also shown as mean fold change of flux following EG00229 or VEGF164 treatment in comparison to the baseline. Asterisks indicate statistical significance (Student’s t-test): **, P < 0.01; ***, P < 0.001; ns, not significant (P > 0.05). (**D**) EG00229 activates p38 and SFK in human ECs. Confluent hCMEC/D3 were treated with EG00229 for the indicated times. Cell lysates were used for immunoblotting with the indicated antibodies, followed by quantification of pixel intensities for P-P38 and P-SFK relative to total P38 and SRC pixel intensities, respectively. GAPDH was used as a loading control. Data are shown as mean fold change ± SD. P38 and SFK phosphorylation levels after EG0229 treatment were quantified at different time points relative to 0 min. Asterisks indicate significant P-values for phosphorylation induction after EG00229 treatment (one-way ANOVA): *, P < 0.05; **, P < 0.01; ns, not significant (P > 0.05).

**Figure S3.**
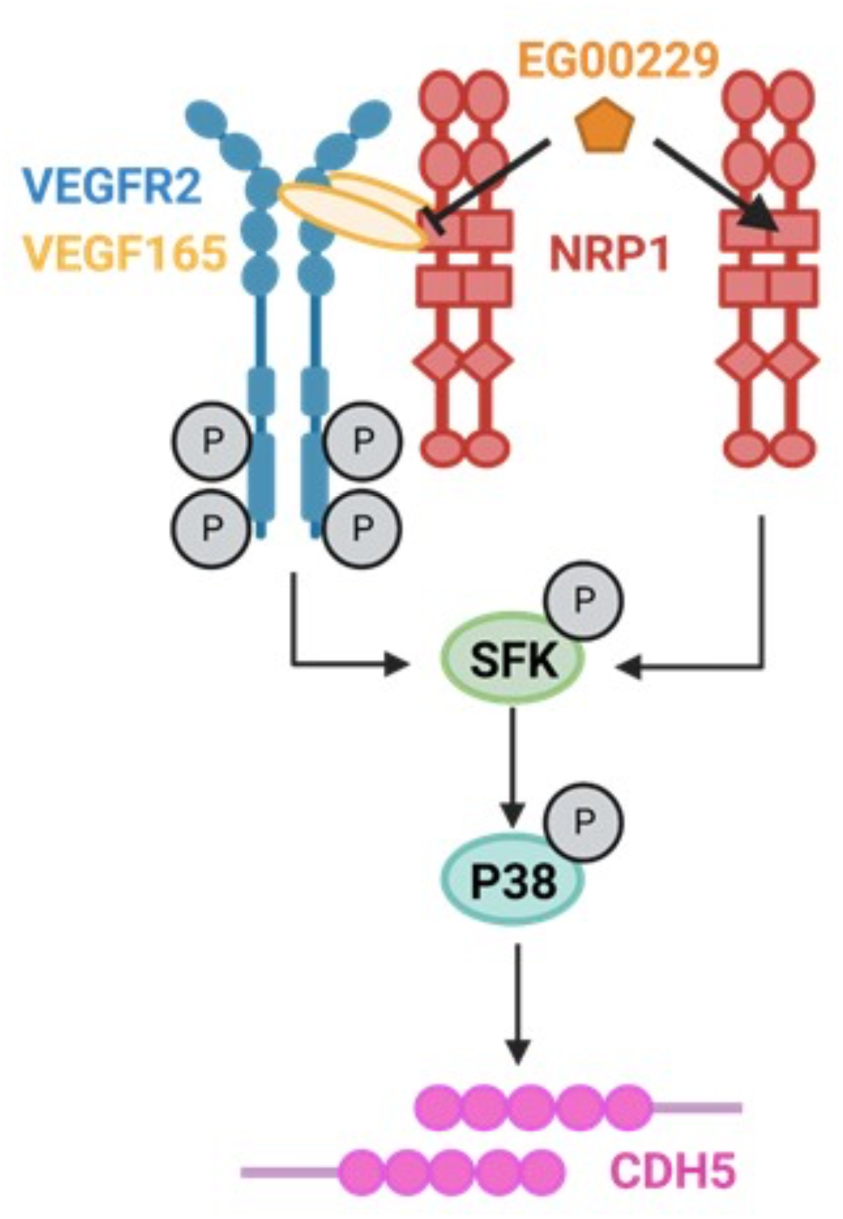
Working model for VEGF164- vs. EG00229-induced signalling in brain and retina ECs. VEGF164 binds to VEGFR2 and NRP1 to induce complex formation, whereas EG00229 binds to NRP1 alone and impairs its interaction with VEGF164. VEGF164 or EG00229 binding to their receptors activates SFK and downstream p38 signalling, which in turn induces CDH5 rearrangements and opening of adherens junctions. Created with BioRender.

